# Reduced antibiotic effect of ciprofloxacin on bacteria in the presence of montmorillonite

**DOI:** 10.64898/2026.05.12.724598

**Authors:** Katharina Axtmann, Benjamin Justus Heyde, Silas Brinkmann, Annette Siskowski, Harald Färber, Lena Marie Juraschek, Melanie Braun, Jan Siemens, Gabriele Bierbaum

## Abstract

Antibiotic residues exceeding selective concentrations for antibiotic-resistant bacteria have been detected in various environments, including manure, wastewater, and effluents from wastewater treatment plants. When these residues come into contact with soils, for instance, due to wastewater irrigation or fertilization with manure, they interact with soil constituents. Soil colloids (1-1000 nm), such as montmorillonite, have been observed to adsorb pharmaceuticals, including antibiotics. We investigated the effect of colloids on the bioavailability of ciprofloxacin and found, that added to bacterial growth medium, montmorillonite reduces, but does not completely prevent, the growth-inhibitory effect of the antibiotic. The bacteria were able to grow at up to roughly double the concentration of ciprofloxacin in the presence of montmorillonite. We show that the incomplete deactivation of ciprofloxacin was most probably caused by medium components that decreased the adsorption of ciprofloxacin to montmorillonite. We conclude that a selective potential of this highly active antibiotic in contaminated soils, which also contain nutrients enabling bacterial growth, cannot be ruled out.

**Environmental implication:** Antibiotics such as ciprofloxacin are frequently detected in water bodies and soils due to wastewater irrigation or manure application. These residues raise concerns about environmental toxicity and antibiotic resistance. This study demonstrates that montmorillonite, a common clay mineral in soils, significantly reduces the antimicrobial efficacy of environmental ciprofloxacin concentrations by sorption. The findings reveal a natural attenuation mechanism that may influence the environmental fate and bioavailability of antibiotics. Understanding such interactions is critical for predicting antibiotic behavior in terrestrial systems and for designing more accurate environmental risk assessments.

## 1 Introduction

Antibiotics are one of the major achievements of modern medicine with a huge impact on human life and health. But ever since the discovery of penicillin in 1928 [1] and its introduction in the 1940s, antibiotic-resistant bacteria (ARB) are an emerging concern [2,3]. In 2021, 4.71 million deaths worldwide were associated with ARB, including 1.14 million deaths attributable to ARB [4]. Forecasts estimate these numbers to increase to 1.91 million deaths attributable to and 8.22 million deaths associated with ARB in 2050 [4] emphasizing the threat to the health system posed by ARB.

High concentrations of antibiotic residues can be found in certain hotspots such as hospital sewage systems and effluents of antibiotic production sites, exceeding Minimal Selective Concentrations (MSCs) and even Minimal Inhibitory Concentrations (MICs), leading to selection of ARB [5,6]. In wastewater and especially wastewater treatment plants (WWTPs), transfer of antibiotic resistance genes among bacteria was shown to be high due to the co-occurrence of different antibiotics and ARB, highlighting the importance of wastewater and WWTPs for the spread of ARB into the environment [7]. On a global scale, most WWTPs are designed to remove solids, nutrients (mainly N and P), and dissolved solids from the wastewater. In contrast, their ability to remove organic micropollutants such as antibiotics or disinfectants is significantly limited. As a result, pharmaceutical residues, including antibiotics, in WWTP effluents are released into the environment [8,9]. Especially in regions where treated or untreated wastewater is used for crop irrigation, both antibiotic residues and ARB come into contact with soils and crops and pose a potential risk for field workers and consumers of the crops [10–12].

Entering soils via wastewater irrigation or manure application, antibiotics and other pharmaceutical residues start interacting with different soil compounds as well as with the soil microbiome [13]. Soils represent a highly complex environment, as their specific composition can be very heterogeneous and may vary a lot, not only between different soil groups but also within one field. Typically, the distribution of chemical substances between the solution phase and the solid phase is described by means of sorption coefficients, such as the distribution coefficient *K*_d_, (*K*_d_ = concentration of chemical sorbed to solid phase/concentration of chemical substance in water). The sorption reduces the mobility of dissolved contaminants in soil, since their concentrations in the mobile phase is reduced. Fluoroquinolones typically exhibit relatively high *K*_d_ values depending on their charge as a function of pH. For example, Nawora et al. (1997) [14] reported a *K*_d_ value of 427 L/kg ciprofloxacin (CIP) to a soil with only 2.5% clay, indicating that the concentration of CIP in free water is only a 1/427 of that sorbed to the soil particles. Consequently, this compound can be considered to be adsorbed almost completely [15]. The sorption of CIP depends on its charge as a function of pH and is controlled mainly by binding of the cationic species to negatively charged clay minerals [16].

Soil colloids form the small particulate phase of soils with particles ranging from 1-1000 nm in diameter [17]. Such colloids often include organic parts (humic substances, bacteria and their extracellular polysaccharides, and decomposed plant material) but also mineral parts such as clay minerals [18,19]. Due to their high surface area and reactivity, clay mineral particles can act as carriers enhancing the mobility of contaminants including antibiotics in soils [20–24]. Montmorillonite (MONT) is one of the most common colloids in clayey soils and is also present in groundwater, underscoring its high mobility [25]. MONT is a member of the smectite group of swelling clay colloids, which have a 2:1-layered structure that allows for interlayer adsorption [26]. Due to their high cation exchange capacity (CEC) and swelling capabilities, smectite-type clays like MONT demonstrate higher sorption capacities than other clay minerals colloids [22,27]. Li et al. (2010) [28] demonstrated the intercalation of tetracycline antibiotics into the interlayers of MONT. Since the interlayer spacing of only a few nanometers prevents access of bacteria, intercalation of antibiotic molecules into interlayers can be expected to reduce the available antibiotic concentrations. A reduced bioavailability and effect of quaternary alkylammonium disinfectants due to their interaction with smectite clay minerals has been demonstrated by Heyde et al. (2020) [29].

Fluoroquinolones are synthetic antibiotics that, due to their high efficacy, are administered in relatively low concentrations to treat bacterial infections in humans and animals. CIP was still the most commonly prescribed fluoroquinolone in Europe in 2022 [30], and it is also the primary metabolite of the frequently used veterinary drug enrofloxacin [31]. Approximately 65% of the administered CIP is excreted in the urine [32], and, therefore, is present in the wastewater system and manure [5,33–35]. Due to their synthetic production, fluoroquinolones are relatively difficult to remove from wastewater for WWTPs by biodegradation, but undergo sorption processes in sewage sludge [36]. Therefore, fluoroquinolones are frequently detected in the effluents of WWTPs and surface waters, resulting in a ubiquitous presence of these antibiotic residues in the environment [33,37–39]. For example, long-term irrigation with wastewater in the Mezquital Valley in Mexico has led to the accumulation of 1.4 μg/kg of fluoroquinolones in the bulk soil [40] and extreme accumulations of up to 26 μg/g of dry weight have been found in the soil of dumpsites that receive, among others, waste of pharmaceutical companies [41]. The MSC of CIP, i.e., the minimum concentration that leads to selection of resistant strains, was estimated to be 17 μg/L for the typical soil bacterium *Acinetobacter baylyi* [5] and 5-10 μg/L for *E. coli* or 1 μg/L for mobile resistance determinants [42]. Furthermore, even estimations of relatively low CIP concentrations in bulk soil may be rather misleading, because CIP mainly adsorbs to the soil in and around the water flow paths. This results in a very uneven distribution of the antibiotic in the soil [43]. Additionally, bacteria interact with colloids [44]. The bacterial surface carries negative charges and binds polyvalent cations, which in turn enable the binding of clay particles that could provide very intimate contact between bacteria and high concentrations of sorbed antibiotics [45,46].

While many studies have been published regarding the sorption of antibiotics to soil colloids, only a limited number of these studies has examined the effect of the colloid-antibiotic-interaction on bacterial growth. This study was therefore designed to measure bacterial growth in the presence of MONT and different concentrations of selected long-lived antibiotics. As bacterial growth is only affected around the minimal inhibitory concentration, diverse bacterial strains with different levels of susceptibility were employed to analyze a wider concentration range. Comparing the concentration available in the truly dissolved fraction of the growth medium with the effect on bacterial growth confirmed that, although CIP is adsorbed by MONT, it still exerts some antibacterial effects. The strength of these effects is shaped by the competitions between nutrients and antibiotic for colloid binding sites.

## 2 Materials and methods

### 2.1 Montmorillonite

The clay mineral used in the experiments was a 2:1 – layer silicate montmorillonite MX-80 (AMCOL, Cheshire, England), which was also studied with respect to its effect on the bioavailability of quaternary alkylammonium compounds (Heyde et al., 2020). The CEC was 106.54 cmolc/kg. The Brunauer–Emmett–Teller specific surface area as determined via N_2_-physisorption-isotherm was 42 m^2^/g.

For the clay purification process, a 1 N sodium acetate buffer (CH_3_COONa, ≥ 99.5%, Merck; CH_3_COOH 100%, Merck), sodium carbonate solution 0.125 g/L (Na_2_CO_3_, 99.9%, Merck), H_2_O_2_ (30%, Merck) and 1 M NaOH (Titrisol, Merck) were used. For the CEC determination, triethylenetetramine (≥ 97%, Sigma-Aldrich) and dehydrated copper (II) sulfate pentahydrate (Ph Eur, Merck) were used in order to prepare a 0.01 mol/L Cu-triethylenetetramine color complex solution.

### 2.2 Minimum inhibitory concentrations

The determination of the MICs followed the guidelines for broth microdilution method of the Clinical and Laboratory Standards Institute (CLSI) in 96-well polystyrene round-bottomed microplates (Greiner Bio-One International GmbH, Kremsmünster, Austria) in Mueller-Hinton broth (MHB, Oxoid Limited, Basingstoke, United Kingdom) with a final concentration of 5 × 10^5^ cells per mL. For *Escherichia coli* ATCC 25922, *Staphylococcus aureus* ATCC 29213, *Enterococcus faecalis* ATCC 29212 and *Bacillus subtilis* W23 the plates were incubated for 24 hours at 37 °C and 170 rpm. For *Acinetobacter baylyi* BD413, the incubation was conducted for 24 hours at 30 °C and 180 rpm. For each strain and antibiotic, the MIC were determined in biological triplicates.

### 2.3 Checkerboards

The bacterial growth response to MONT and the potential buffering effect of the colloid on the susceptibility of bacterial cultures to the exposure to the tested antibiotics (CIP, sulfamethoxazole (SMX), clindamycin (CLI) and erythromycin (ERY)) were tested in checkerboard assays. The maximum concentrations of colloids were adjusted in such a way, that turbidity measurements of bacterial growth were still possible at the highest concentration of MONT (256 μg/mL).

All experiments were performed in a total volume of 200 μl in 96-well round-bottomed polystyrene microplates with sterile transparent plastic lids (Greiner Bio-One International GmbH, Kremsmünster, Austria). The concentrations of MONT tested ranged from 0 to 256 μg/mL, and the antibiotic concentrations tested were around the MIC of the respective strain. An inoculum of the respective bacterial strain was prepared from an overnight preculture and added to each well, with the aim of achieving a final concentration of 5 × 10^5^ cells per mL. The plates were then shaken in a microplate reader (TECAN Infinite® 200 PRO) for 20 hours at 30 °C for *A. baylyi* BD413 or at 37 °C for all other strains. The optical density at 600 nm (OD_600_) was automatically determined every 15 minutes. For each strain and antibiotic tested, a minimum of three biological replicates were performed.

### 2.4 Sorption of ciprofloxacin and sulfamethoxazole to montmorillonite

To assess the sorption capacity of the MONT-CIP combination, 20 ng/mL CIP or SMX was added to 40 mL sterile deionized water containing 640 ng/mL MONT, since measurements in MHB were not possible. For each antibiotic, three technical replicates were carried out. For the extraction of the truly dissolved fraction, 30 mL of the liquid sample were transferred to ultracentrifuge tubes and then filled with Millipore to the maximum filling volume of 66 mL. The samples were then centrifuged at 45,000 rpm for 26 hours. The resulting supernatant after ultracentrifugation represented the truly dissolved fraction and the remaining pellet the colloid fraction. Subsequently, 50 mL of the supernatant was transferred and completely evaporated. The sample was then re-dissolved in methanol and passed through a filter (0.22 μm). A further complete evaporation and re-dissolution of the sample was carried out with 900 μL H_2_O/ MeOH (90:10; v:v) and 100 μL internal standard solution in GC vials. The truly dissolved fraction from these suspensions was measured on an Acquity H-class UPLC (Waters, Eschborn, Germany) according to a modified method of Dalkmann et al. (2012) [40] and Kappenberg & Juraschek (2021) [47].

In order to confirm the results from the described sorption experiments, different concentrations of CIP and the MONT in were pre-incubated together in distilled water for 24 hours at room temperature and with horizontal shaking. Subsequently, the colloid was removed by centrifugation (45,000 rpm, 26 hours, 4 °C). The obtained supernatants were filtered (0.22 μm) and mixed 1:1 with two-fold MHB. The suspensions were inoculated with *A. baylyi* BD413 and incubated in 96-well plates as described above. The experiment was performed three times to achieve biological replicates. Controls were performed in MHB.

### 2.4 Effect of montmorillonite on bacterial growth

To evaluate possible growth-inhibitory effects of MONT on bacterial growth in the absence of antibiotics, MHB was pre-incubated with different MONT concentrations for 24 hours at room temperature and horizontal shaking. Subsequently, the colloid was removed by centrifugation (15,000 g, 15 minutes, no brake). Half of the supernatant was mixed in a 1:1 ratio with twofold concentrated MHB to reconstitute growth factors that might have been adsorbed by the colloid. Both the mixed and unmixed supernatants and the non-centrifuged MHB-MONT suspensions were inoculated with the five different strains and incubated in 96-well plates as described above. The experiment was performed three times to achieve biological replicates.

### 2.6 Ion-release of montmorillonite

A total of 50 mL of sterile deionized water was inoculated with 256 μg/mL or 0 μg/mL MONT (each with five replicates) in Falcon tubes for 24 hours at room temperature and horizontal shaking. Subsequently, 2 g of CaCO_3_ were added to each tube to ensure complete removal of the colloid and then shaken for two minutes. The colloid was then pelleted by centrifugation (10,000 g, 20 minutes, no brake) and the supernatant was carefully transferred to a new tube without disturbing the pellet. The ion concentrations within the supernatants were then measured by ICP-MS.

To ascertain the effect of Al^3+^ and Fe^2+^ ions released by MONT on bacterial growth dynamics, solid AlCl_3_ and FeSO_4_ were exposed to UV radiation for 30 minutes and then dissolved in sterile deionized water. In a checkerboard assay, serial dilutions of both metals were prepared in 96-well plates within MHB. The highest concentrations corresponded to the double of the mean values previously determined for Fe and Al ion from the MONT supernatant (18,830 μg/L or 2,862 μg/L respectively). The plates were then inoculated with the five different strains and incubated in 96-well plates as previously described. The experiment was carried out three times to achieve biological replicates.

### 2.7 Statistical analysis

To statistically assess the effects of different combinations of MONT and antibiotics on bacterial growth, a two-way ANOVA model was fitted using the *aov()* function in R, with concentrations of the antibiotic and concentrations of MONT as categorical factors. Estimated marginal means (EMMs) were computed for each combination of antibiotic and MONT using the *emmeans* package [48]. Custom pairwise comparisons between specific concentration combinations were performed via contrast vectors applied to the EMMs. This allowed the estimation of differences in mean relative growth between selected concentration pairs while accounting for interactions. Statistical significance of contrasts was evaluated based on the corresponding p-values.

## 3. Results

### 3.1 Minimum inhibitory concentrations of tested antibiotics

MICs of the five indicator strains for four long-lived antibiotics that might accumulate in soil due to their slow dissipation are shown in Table 1. The indicator strain panel comprises pathogenic and soil bacteria with a wide range of MICs against the tested antibiotics.

**Table 1:** The MICs of *A. baylyi* BD413, *E. coli* ATCC 25922, *S. aureus* ATCC 29213, *E. faecalis* ATCC 29212 and *B. subtilis* W23 against CIP, clindamycin (CLI), SMX and erythromycin (ERY). The values were determined using the EUCAST method. n.d.= not determined.

| MIC [$\mu\text{g/mL}$] (24 h) | CIP | CLI | SMX | ERY |
| --- | --- | --- | --- | --- |
| *A. baylyi* BD413 | 0.05 | 2 | 2 | 1 |
| *E. coli* ATCC 25922 | 0.01 | 128 | 8 | 64 |
| *S. aureus* ATCC 29213 | 0.3 | 16 | 4 | 1 |
| *E. faecalis* ATCC 29212 | 0.5 | 0.25 | >256 | 2 |
| *B. subtilis* W23 | 0.03 | n.d. | n.d. | n.d. |

### 3.2 Montmorillonite enables growth in the presence of ciprofloxacin

The OD_600_ of *A. baylyi* BD413, *E. coli* ATCC 25922, *S. aureus* ATCC 29213, *E. faecalis* ATCC 29212 and *B. subtilis* W23 was measured in the presence of various concentrations of CIP and MONT during 20 hours of incubation. The results, expressed as the relative growth (growth compared to growth without antibiotic and colloid in percent) for each strain and combination, are shown in Figure 1. Standard deviations are listed in Figure S1 of the supplemental data. The presence of the highest MONT concentration resulted in growth at higher CIP concentrations for all five strains. At 60 μg/L CIP no growth was observed for *A. baylyi* BD413 without MONT, while in combination with 256 μg/mL MONT increased growth equaling 44% of the growth in the absence of CIP was observed (p < 0.001). Similar results were seen for *E. coli* ATCC 25922. At the respective CIP MIC of 10 μg/L, 56% relative growth was observed in the presence of 256 μg/mL MONT, while no growth was detected at lower colloid concentrations (p < 0.001). For *S. aureus* ATCC 29213 and *E. faecalis* ATCC 29212, approximately 30% relative growth was measured in the presence of 256 μg/mL MONT at the respective CIP MIC in MONT-free MHB (Figure 1). Even though, the relative growth at this colloid concentration was lower than for the two Gram-negative strains tested, the observed growth was still significantly stronger in the presence of MONT compared to the growth at the same CIP concentration without MONT (p < 0.001). *B. subtilis* W23 demonstrated 100% relative growth at the MIC for the three highest MONT concentrations and 35% relative growth at 2 x MIC in the presence of 256 μg/mL MONT (p < 0.001 compared to the same CIP concentrations without MONT). The other antibiotics tested (SMX, CLI, and ERY), resulted no growth above the respective MIC in the presence of MONT for the five bacterial strains tested. The results for these antibiotics for *A. baylyi* BD413 are shown in Figure S2-S4 of the supplemental material.

**Figure 1:**
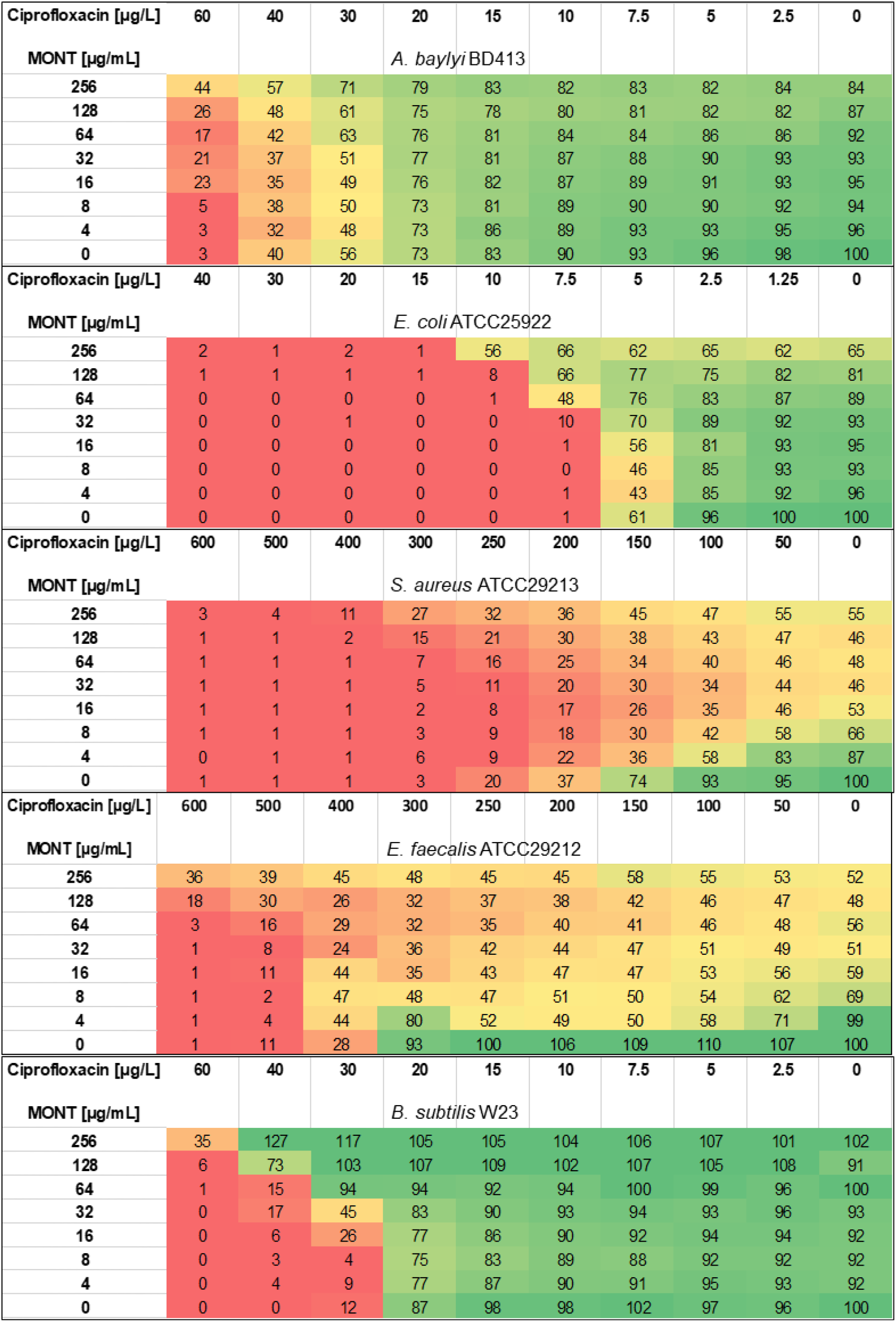
CIP-MONT-Checkerboards. Relative growth [%] of *A. baylyi* BD413, *E. coli* ATCC 25922, *S. aureus* ATCC 29213, *E. faecalis* ATCC 29212 and *B. subtilis* W23 with different concentrations of CIP and MONT in a 96-well plate. The optical density after 20 h of incubation in each well was compared to growth without antibiotic and colloid and expressed as percentage. Standard deviations are listed in Figure S1. Due to the low MICs obtained with CIP, concentrations are indicated as μg/L.

### 3.3 Sorption of ciprofloxacin to montmorillonite

To confirm CIP sorption to MONT as main driver of sustained bacterial growth above MICs, suspensions of the colloid in distilled water with CIP or SMX were subjected to ultracentrifugation and the supernatants analyzed by UPLC. The concentrations of the two antibiotics, which were measured in the supernatant, the truly dissolved concentrations, are listed in Table 2. Of the initial CIP concentration, 0.29% was detected within the truly dissolved phase. In contrast, 55.15% of the initial SMX concentrations were measured within the truly dissolved phase.

**Table 2:** In 40 mL Milli-Q water, 20 ng/mL of CIP or SMX were mixed with 640 ng/mL MONT. The truly dissolved fractions were measured on an Acquity H-class UPLC.

|  | Initial concentration<br>[ng/mL] | Concentration in the truly dissolved phase<br>[ng/mL] |
| --- | --- | --- |
| CIP | 20 | 0.06 ± 0.01 (0.3%) |
| SMX | 20 | 11.0 ± 0.79 (55.2%) |

To confirm this strong sorption of CIP observed in distilled water by a growth experiment, different concentrations of CIP and the MONT were incubated for 24 hours in distilled water and then ultra-centrifuged to remove the colloid with potentially sorbed CIP. The supernatants were mixed with twofold MHB and then used for growth experiments with *A. baylyi* BD413. Full growth was observed for all CIP concentrations ranging around the MIC even in the presence of the lowest tested MONT concentration (4 μg/mL). Even at the tenfold CIP-MIC, 91% relative growth was observed at 256 μg/mL MONT. Conversely, no growth was observed at CIP concentrations above the MIC in the absence of MONT in controls (Figure 2), demonstrating that the CIP had indeed been removed together with the colloids. However, growth inhibition was strongly increased, when the sorption was performed in MHB instead of distilled water. Standard deviations are listed in Figure S5.

**Figure 2:**
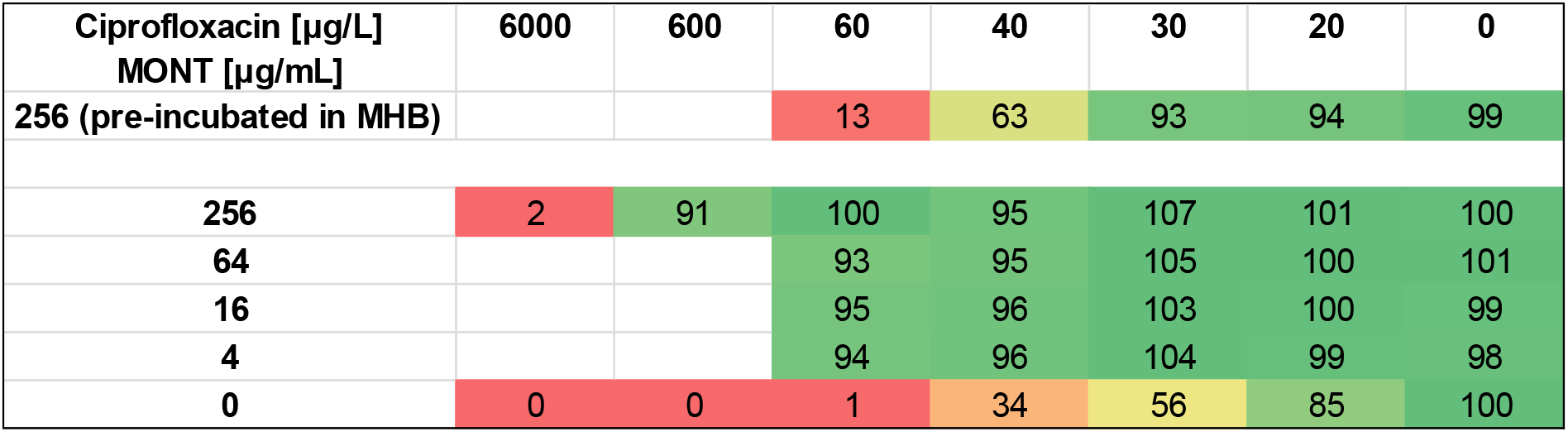
CIP-MONT-Sorption. Relative growth [%] of *A. baylyi* BD413 in the supernatants of pre-incubated suspensions of CIP and MONT in distilled water or MHB (only first row). After 24 hours of pre-incubation, the colloid was removed via ultra-centrifugation and the supernatants mixed with twofold MH before inoculation of bacteria. Growth was measured for 20 hours at 30 °C with horizontal shaking, then final OD_600_-values were compared to MHB cultures without antibiotic and colloid. Standard deviations are listed in Figure S5.

### 3.4 Montmorillonite inhibits growth of *S. aureus* and *E. faecalis*

Reduced growth of both Gram-positive cocci and to a lesser extent *E. coli* ATCC-25922 in the presence of MONT and absence of CIP indicated a growth inhibitory effect of MONT (p < 0.0001). Whereas the growth of *B. subtilis* W23 and *A. baylyi* BD413 was not or only slightly impeded by the colloid, *E. coli* ATCC 25922 showed a relative growth of 65% and *S. aureus* ATCC 29213 and *E. faecalis* ATCC 20212 exhibited a final OD_600_ of 55% and 52% in the presence of the highest colloid concentration and in absence of the antibiotic compared to their growth in colloid-free MHB. The growth inhibitory effect of MONT can be related to different potential processes. Morrison et al. (2016) [49] demonstrated production of the toxic H_2_O_2_ during the first 24 h after mixing of medium with Oregon blue medicinal clay, in addition, a release of iron and aluminum ions from the colloids was found to exert bactericidal effects. Furthermore, MONT might sequester essential nutrients required by the bacteria from the culture medium, since *S. aureus* and *E. faecalis*, in contrast to *A. baylyi, E. coli* and *B. subtilis*, are fastidious organisms and have to take up vitamins of the B group [50,51]. To further elucidate the growth inhibitory effect of the colloid, MHB was mixed with different concentrations of MONT. These mixtures were either used at once (“fresh”) or after a pre-incubation for 24 h (“24 h”), after which the colloid was removed by centrifugation. Half of the supernatant was then mixed with twofold concentrated MHB to reconstitute any essential nutrients that might have been adsorbed by the colloid (“SN+2xMH”), the rest was tested without addition (“SN”). The relative growth values of the five strains grown in both, the mixed “SN+2xMH” and unmixed supernatants (“SN”), and in the non-centrifuged MHB-MONT suspensions (“fresh” and “24 h”) are shown in Figure 3. Standard deviations are listed in Figure S6. The pH values of the colloid suspension were measured and compared to the pH values of the growth controls without MONT. No significant pH differences between the different strains or after colloid addition were observed.

**Figure 3:**
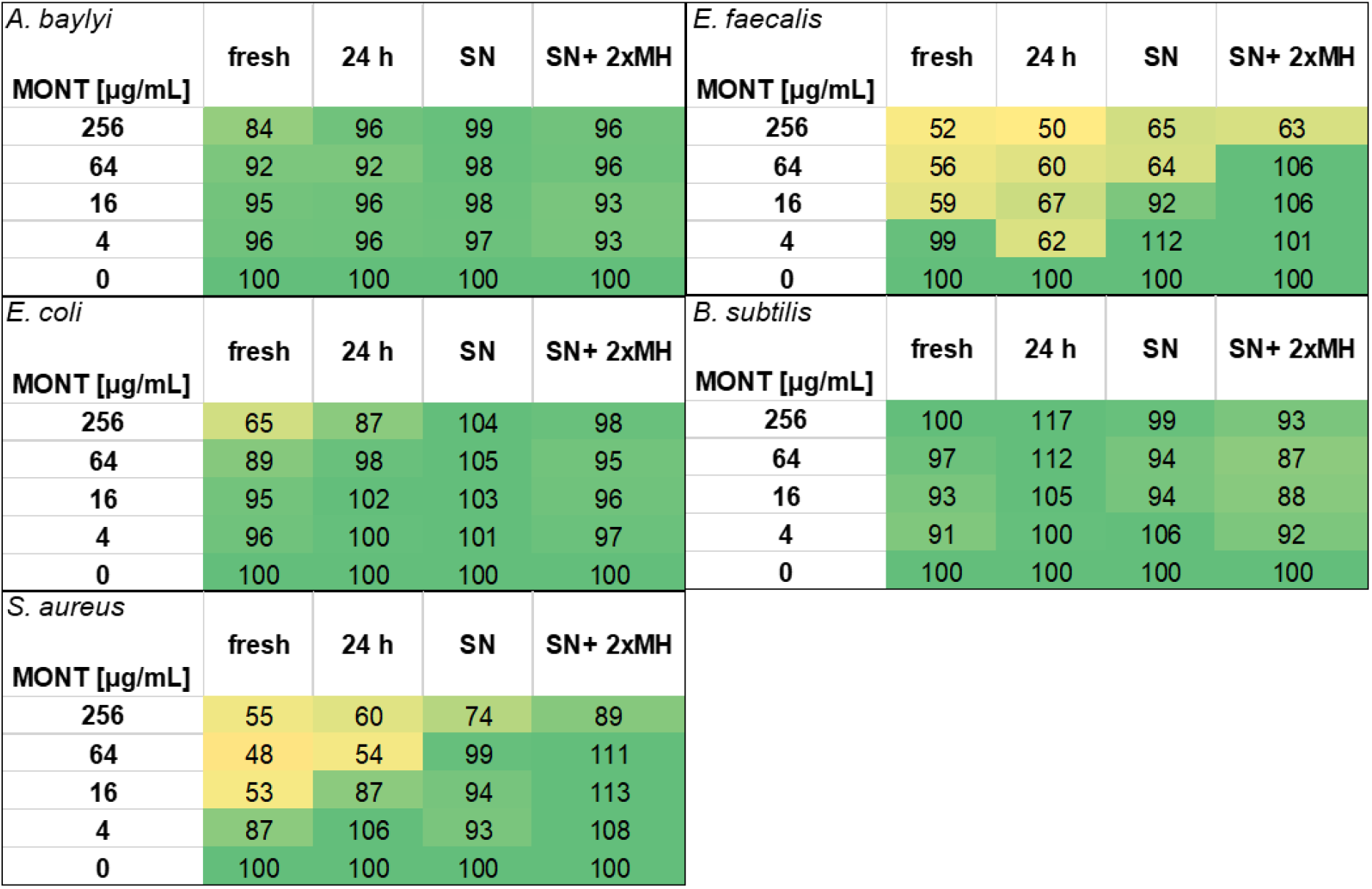
Growth inhibitory effect of MONT. Relative growth [%] after 20 h of incubation of *A. baylyi* BD413, *E. coli* ATCC 25922, *S. aureus* ATCC 29213, *E. faecalis* ATCC 29212 and *B. subtilis* W23 in a suspension of MHB with freshly added MONT (fresh), suspensions of MHB and MONT pre-incubated 24 h before inoculation (24 h), the supernatant of the 24 h pre-incubated suspension after removal of colloids (SN) and the same supernatant mixed with twofold concentrated MHB to reconstitute nutrients adsorbed by MONT (SN+ 2xMH). Growth values were compared to growth without colloid. Standard deviations are listed in Figure S6 of the supplemental.

For *A. baylyi* BD413 and *B. subtilis* W23, no discernible growth inhibition was observed after 24 h in all samples. In contrast, the relative growth of *E. coli* ATCC 25922 at the highest MONT concentration was significantly higher (87%) in the pre-incubated suspension “24 h” than in the freshly mixed (65%) medium, and increased even further to 104% within the supernatant (“SN”) (p < 0.001). In addition, no discernible growth inhibition was observed within the supernatants reconstituted with MHB (“SN+2xMH”).

*S. aureus* ATCC 29213 and *E. faecalis* ATCC 29212 demonstrated impaired growth in the fresh and pre-incubated MONT-MHB suspension (“24 h”) (p < 0.001 for comparison of 256 μg/mL to 0 μg/mL MONT). Following the removal of the colloid, *S. aureus* ATCC 29213 exhibited a reduced growth value (74%) in the supernatant (“SN”) that had been pre-incubated with the highest MONT concentration (256 μg/mL). The addition of twofold concentrated MHB (“SN+2xMH”) resulted in an enhancement of relative growth to 89% (p < 0.05 for comparison with 24 h pre-incubated MONT).

*E. faecalis* ATCC 29212 demonstrated a relative growth of 65% in supernatant that had been pre-incubated with the two highest MONT concentrations (p < 0.05). Mixing the supernatant with twofold concentrated MHB (“SN+2xMH”) led to an augmentation in relative growth at 64 μg/mL of the colloid (p < 0.001), but this effect was not observed for the highest concentration of 256 μg/mL. These data indicated that at least a part of the growth inhibition seen with the cocci might be caused by sequestration of essential nutrients by the colloids, since these two species show a higher dependence on essential nutrients (vitamins, amino acids) than the other three strains [50].

To evaluate a potential effect of ions, esp. Al and Fe, released by MONT into liquid environments on bacterial growth, Milli-Q water was incubated with the colloid for 24 hours. After removal of the colloid with CaCO_3_ and centrifugation, the ion concentrations in the supernatant were determined by ICP-MS. The results of this analysis are listed in Table 3.

**Table 3:** Ion-concentrations in Milli-Q water incubated for 24 hours with 0 μg/mL or 256 μg/mL MONT measured via ICP-MS. After incubation, the colloid was removed by adding CaCO_3_ followed by centrifugation. Concentrations were determined within the supernatants.

| µg/L | Al | Fe | Pb | U | Si | Ba | Th | Ti | Na | Ca | Mg |
| --- | --- | --- | --- | --- | --- | --- | --- | --- | --- | --- | --- |
| **water** | <10 | <10 | <1 | <0.1 | <0.1 | <1 | <0.1 | <1 | 8720<br>(±449.0) | 11440<br>(±155.3) | <100 |
| **water**<br>**+**<br>**MONT** | 9,415<br>(±755.8) | 1,431<br>(±126.3) | 3<br>(±0.3) | 1<br>(±) | 9,700<br>(±845.6) | 9<br>(±0.4) | 4<br>(±0.2) | 52<br>(±3.2) | 17,000<br>(±0.0) | 27,000<br>(±2915.5) | 1,700<br>(±122.5) |

## 4. Discussion

### 4.1 Sorption of CIP to MONT

The sorption of several antibiotics to soils and soil colloids, such as MONT, has been demonstrated in several previous studies [22, 52–55]. In particular, Chen et al. (2015) [52] reported a maximum CIP sorption of 23 g/kg for MONT, which was attributed to the high CEC of the colloid (990 mmol/kg) [22]. Furthermore, Wang et al. (2011) showed that CIP can sorb to both the inner and outer surfaces of MONT [53]. Carrasquillo et al. (2008) [22] described similar results and stated that the CIP sorbs to MONT by its cationic amine into the MONT interlayers through cation exchange and via electrostatic forces between the MONT surface and the aromatic ring of the antibiotic [22]. Interestingly, a high complexation ability of CIP to Fe and Al was also described [56,57]. Since the MONT tested here releases high concentrations of both elements into the supernatant (see Table 3), a reduced bioavailability of the antibiotic due to complexation cannot be excluded. The strong sorption of CIP to MONT was also confirmed by our results, as less than 1% of CIP was found in the truly dissolved fraction, when adsorption was tested in distilled water. The consistent strong sorption of CIP to soil mineral colloids underscores the potential impact of colloid-antibiotic interactions on bacterial growth and enhances the potential loss of effectiveness of CIP when colloids are abundant.

### 4.2 Reduced bioavailability and efficacy of CIP due to MONT

As illustrated in Figure 1, the growth of all five strains significantly (p < 0.001) exceeded their respective CIP MIC at the highest MONT concentration, suggesting that the colloid reduces the bioavailability of the antibiotic. It is noteworthy that the ratio of CIP and MONT used in the adsorption experiment corresponds to the combination of 8 μg/L CIP and 256 μg/mL MONT in the checkerboard experiments shown in Figure 1. This concentration was tested with *E. coli* ATCC 25922, which was the only bacterium with an MIC of 7.5 μg/L. In the presence of MONT, growth at 7.5 μg/L was as strong as in the control without antibiotic, but at 15 μg/L growth was inhibited also in the presence of MONT (see Figure 1).

Therefore, the growth experiments indicate that despite the sorption of CIP to MONT (maximum CIP sorption for MONT of 23 g/kg reported by Chen et al. (2015) [52], about 50 % of the adsorbed CIP is still available for the bacteria. For example, at a CIP concentration of 600 μg/L in combination with 256 μg/mL MONT, the ratio of antibiotic to colloid is at 2.34 g/kg. In spite of this, a growth inhibition was observed for *E. faecalis* ATCC 29212 (36% relative growth compared to 52% in the absence of CIP (p<0.05) and for *S. aureus* ATCC 29213 (no growth, compared to 55% in the absence of CIP (p < 0.001). However, full growth would have been expected after complete sorption of CIP by the colloid. The reduced yet still detectable effect of CIP in our experimental model could be attributed to either unsorbed or detached molecules of the antibiotic or externally sorbed CIP, which retained its antibiotic effect on the tested bacterial strains. Okaikue-Woodi et al. (2018) [58] reported a higher degree of fluoroquinolone sorption to MONT at acidic pH compared to neutral pH. CIP sorption to smectite-containing Mexican soils decreased with increasing pH due to a decreasing fraction of the cationic species [59]. The growth experiments were performed here in MHB with a pH of 7.3 in deionized water according to the manufacturer’s information (http://www.oxoid.com/UK/blue/prod_detail/prod_detail.asp?pr=CM0405, accessed on March 5^th^ 2025).

Therefore, the maximum sorption may not have been reached due to the neutral pH or the presence of media components competing with MONT for binding sites of the colloid (Figure 2). The latter hypothesis is supported by experiments that tested first adsorption of CIP to MONT and after removal of the loaded colloid, growth of *A. baylyi*. Here full growth was obtained with high CIP concentrations. However, this effect was observed only, if adsorption was performed in distilled water and growth decreased if adsorption was performed in the presence of the growth medium (p < 0.01) (Figure 2). Performing all experiments in distilled water would not be possible, as bacterial growth, which is necessary for selection, requires nutrients [60]. Additionally, as discussed below, the growth of *S. aureus* and *E. coli* was inhibited by nutrient sequestration in the presence of MONT, confirming the interaction of MONT and media components (Figure 3). On the other hand, it is well possible, that - in the presence of CIP loaded MONT - bacteria may bind to or interact with the colloid particles and are able to release further CIP molecules bound on the surface of the particles, resulting in an inhibitory effect. Bacterial metabolites that increase the pH in the vicinity of bacteria associated with colloids, such as amines after decarboxylation of amino acids, might also play a role in the remobilization of colloid-bound antibiotics. This suggests that CIP bound to soil colloids may still exert selective pressure on bacteria. Chen et al. (2015) [52] observed an inhibition of *E. coli* DH5α by CIP-loaded MONT compared to a MONT suspension devoid of CIP. However, a much higher CIP concentration of 1 μg/mL in combination with 450 μg/mL MONT was used in their experiments [52]. In our experiments, we determined an MIC for *E. coli* ATCC25922 against CIP of 0.01 μg/mL, a value that is consistent with the established distribution parameters set forth by the European Committee on Antimicrobial Susceptibility Testing (EUCAST) (https://mic.eucast.org/search/, accessed on March 5^th^ 2025). Hence, the CIP concentrations used in our experiments were approximately 100 times lower than those reported by Chen et al. (2015) [52] in combination with a relatively much higher MONT concentration. Our results regarding the deactivation of CIP due to sorption align with results of Rosendahl et al. (2012) [61] illustrating the inactivation of the fluoroquinolone antibiotic difloxacin due to its strong sorption and formation of non-extractable residues. Blau et al. (2017) [62] showed that when applied with manure, antibiotic residues had a stronger impact on soil microbiomes and ARG abundance in sandy soils than in loamy soils. This emphasizes the importance of understanding the effect of soil colloids on antibiotic bioavailability.

### 4.3 Effects of MONT on the bioavailability and efficacy of other antibiotics

For the other antibiotics tested in this study, no growth was detected above the respective MIC (Figures S2-S4). This may be due to the following: (i) the adsorption is lower than that of CIP and (ii) the concentrations needed for growth inhibition are much higher for the other substances than for CIP, meaning that higher antibiotic concentrations had to be added to the experiments and (iii) that these high concentrations may exceed the absorption capacities of the added MONT, which could not be increased proportionally due to its turbidity. Lu et al. (2014) [63] reported less than 5% sorption of SMX on MONT after 5 h of incubation of a solution of 100 μg/mL SMX with 2.5 mg/mL MONT. Gao & Pedersen (2005) [64] reported only low sorption of SMX on MONT over a broad pH range and Avisar et al. (2010) [65] expected sorption to be negligible at ambient pH conditions. Experiments of Dalkmann et al. (2014) [59] showed a much weaker sorption of SMX compared to CIP in smectite-containing Mexican soils.

From the initial concentration of 20 ng/mL SMX, approximately 45% was sorbed to the tested MONT at a concentration of 640 ng/mL (see Table 2), indicating that sorption at low antibiotic concentrations may be higher than reported elsewhere. CLI sorbs to MONT only under acidic pH conditions, favoring the cationic form of the antibiotic ([66], i.e. conditions which would not allow bacterial growth of the strains tested here. Only few studies have looked into the sorption of ERY or other macrolide antibiotics to MONT. While Hanamoto & Ogawa (2019) [67] could show sorption affinities of azithromycin, an ERY derivative, to MONT, Arif et al. (2023) [68] could increase the removal of azithromycin from water by MONT due to attachment of biochar to the surface of the colloid. More research is needed to improve our understanding of the interactions between ERY and colloids and their effect on bacterial growth. A non-optical method that allows for the addition of higher MONT concentrations would have to be established, and indicator strains that are not inhibited by MONT would have to be used.

### 4.4 Effect of MONT on bacterial growth

Only a few studies have reported the antibacterial properties of certain medicinal clays, including montmorillonites [49,69–71]. As illustrated in Figure 1, the MONT used in this study exhibited a significant (p < 0.0001) inhibitory effect on the growth of three of five bacterial strains in the absence of CIP. While Tong et al. (2005) [71] did not observe a bactericidal effect of MONT on *E. coli* K88, Morrison et al. (2016) [49] described 100% killing of various Gram-positive and Gram-negative strains by an illite-smectite clay that contained also pyrite. Noteworthy, the colloid concentrations of 500 mg/mL used by the authors Morrison et al. (2016) [49] were much higher than the highest MONT concentration of 256 μg/mL used in our studies. Comparably high concentrations of a clay composed of 50% Fe-smectite exhibited bactericidal effects against several potentially pathogenic Gram-negative bacteria and resulted in growth reduction of several *S. aureus* strains [69]. In contrast to these results, the highest MONT concentration tested in our study exhibited the strongest growth inhibition of *S. aureus* ATCC 29213 and *E. faecalis* ATCC 29212 (55% and 52% relative growth respectively), compared to *E. coli* ATCC 25922 (65% relative growth). For *S. aureus* ATCC 29213, the effects were nearly totally compensated by addition of twofold MH to the supernatant (p < 0.001), indicating that sorption of essential nutrients was responsible for the observed growth-inhibitory effect of MONT. For *E. faecalis* ATCC 29212, a growth inhibition remained after addition of twofold MHB. Clay suspensions of an illite-smectite were shown to produce hydrogen peroxide (H_2_O_2_) in the presence of pyrite in different media, resulting in a bactericidal effect [49]. However, the presence of pyrite in the MONT used in our experiment is unlikely due its pre-treatment with the strongly oxidizing H_2_O_2_.

Al and Fe have been shown to bind to the membranes of *E. coli* ATCC 25922, where Fe^2+^ can cause ROS stress (see below) and Al^3+^ leads to misfolded outer membrane proteins [49]. Fe^2+^ also enters *E. coli* cells via siderophores [72], high-affinity transporter systems [73] or by a CorA magnesium transporter system due to its similar charge and size to Mg^2+^ [74]. Inside the cell, an overload of Fe^2+^ leads to generation of hydroxyl radicals that react with intracellular molecules [49].

Pre-incubation of MHB with MONT resulted in stronger growth than within freshly mixed MHB with MONT (see Figure 3), which might be explained by reduced H_2_O_2_ production after 24 h of pre-incubation [49]. Removal of the colloid by centrifugation after the pre-incubation with MHB further increased growth, especially visible for *S. aureus* ATCC 29213 and *E. faecalis* ATCC 29212 (p < 0.05), suggesting a direct effect of the colloid on the cells when present in medium. The addition of twofold concentrated MHB to the supernatant after incubation with MONT, restored only 89% of the relative growth of *S. aureus* ATCC 20213 compared to the growth control in MHB. This suggests that during pre-incubation, certain MONT components that inhibited the bacteria, such as metal ions, dissolved into the medium and remained in the supernatant after centrifugation. Morrison et al. (2016) [49] determined the MIC and MBC of a leachate of an illite-smectite clay using a rich medium and a minimal medium. The observed MIC and MBC of *E. coli* ATCC 25922 were six- or three-times higher in rich medium than in the minimal medium. This result was explained by the higher buffer capacity of the rich medium, preventing complete oxidation of the metal ions released from the clay into the leachates.

For further investigation, the ion release of MONT into liquid environments was measured by ICP-MS (see Table 3). The elevated calcium concentrations observed in the water control sample can be attributed to the presence of CaCO_3_, which was added to ensure complete removal of the colloid. Of the elemental concentrations evaluated, Al and Fe were hypothesized to have the most significant influence on bacterial growth. Similar measurements were performed by Morrison et al. (2016) [49] in the leachates of an illite-smectite clay. Per mg MONT we measured 0.1 nM Fe and 1.4 μM Al, while Morrison et al. (2016) [49] measured 81.1 nM Fe (combined values of Fe^2+^ and Fe^3+^) and 27.9 nM Al, indicating different compositions of the clays and a much higher, growth-inhibitory release of ions by the medicinal clay used in the study of [49].

## 5. Conclusion

This study underscores the impact of colloid-antibiotic interactions on antibiotic efficacy. MONT significantly reduces the bioavailability of CIP to bacteria, as this effect was evident in all five bacterial strains tested, with growth observed at CIP concentrations that would otherwise be inhibitory. The strong sorption of CIP to MONT resulted in less than 1% of the antibiotic remaining in the truly dissolved phase in distilled water, supporting the hypothesis that adsorption to the colloid sequesters CIP. In contrast, in growth medium, the MIC of all strains increased only by about one titer step in the presence of MONT, indicating that about 50 % of the added CIP was still bio-available. In addition, MONT also exhibited direct inhibitory effects on some bacterial strains, particularly the two tested Gram-positive cocci, likely due to ion release and nutrient sequestration.

These results underscore the need to consider colloid-mediated sorption when evaluating the environmental fate and microbial impact of antibiotics, especially in soils where MONT or other colloids from the smectite group, are prevalent. In addition, our results raise the question under which conditions CIP is released from colloids, surface or interlayers, and become bioavailable to bacteria.

Future research should focus on the physicochemical factors controlling antibiotic release from colloid surfaces and interlayers, such as pH, competing ions and organic compounds, and microbial activity. A better understanding of these dynamics is essential for evaluating the selective pressure exerted by colloid-bound antibiotics in complex environmental systems and their implications for antimicrobial resistance development. In the end, to conclude about the role of soil colloids for the bioavailability of antibiotics, experiments are needed with real soils that are fertilized with antibiotic containing manure, mirroring the complex interplay of different soil constituents under realistic conditions, also accounting for varying properties of different soils.

## Supporting information

Supplemental Materials includes all data

## Acknowledgements/Funding

This research was funded by the Deutsche Forschungsgemeinschaft (DFG, German Research Foundation) in the framework of the Research Unit FOR 5095: Pollutant – Antibiotic Resistance – Pathogen Interactions in a Changing Wastewater Irrigation System, project number 431531292.

## CRediT authorship contribution statement

**Katharina Axtmann:** Conceptualization, Data curation, Formal analysis, Investigation, Methodology, Writing – original draft, Writing – review & editing. **Benjamin Justus Heyde:** Methodology, Project administration, Writing – original draft, Writing – review & editing. **Silas Brinkmann:** Methodology, Data curation. **Annette Siskowski:** Methodology, Data curation. **Harald Färber:** Methodology, Writing – review & editing. **Lena Marie Juraschek:** Methodology, Writing – original draft. **Melanie Braun:** Writing – review & editing, Supervision, Project administration, Funding acquisition, Conceptualization. **Jan Siemens:** Writing – review & editing, Supervision, Project administration, Funding acquisition, Conceptualization. **Gabriele Bierbaum:** Writing – review & editing, Supervision, Project administration, Funding acquisition, Conceptualization.

## Declaration of Competing Interest

The authors declare no financial interests/personal relationships which may be considered as potential competing interests.

## Data availability

Data will be made available on request.

## Notes

### Competing Interest Statement

The authors have declared no competing interest.

