## Supplemental Materials includes all data for "Reduced antibiotic effect of ciprofloxacin on bacteria in the presence of montmorillonite"

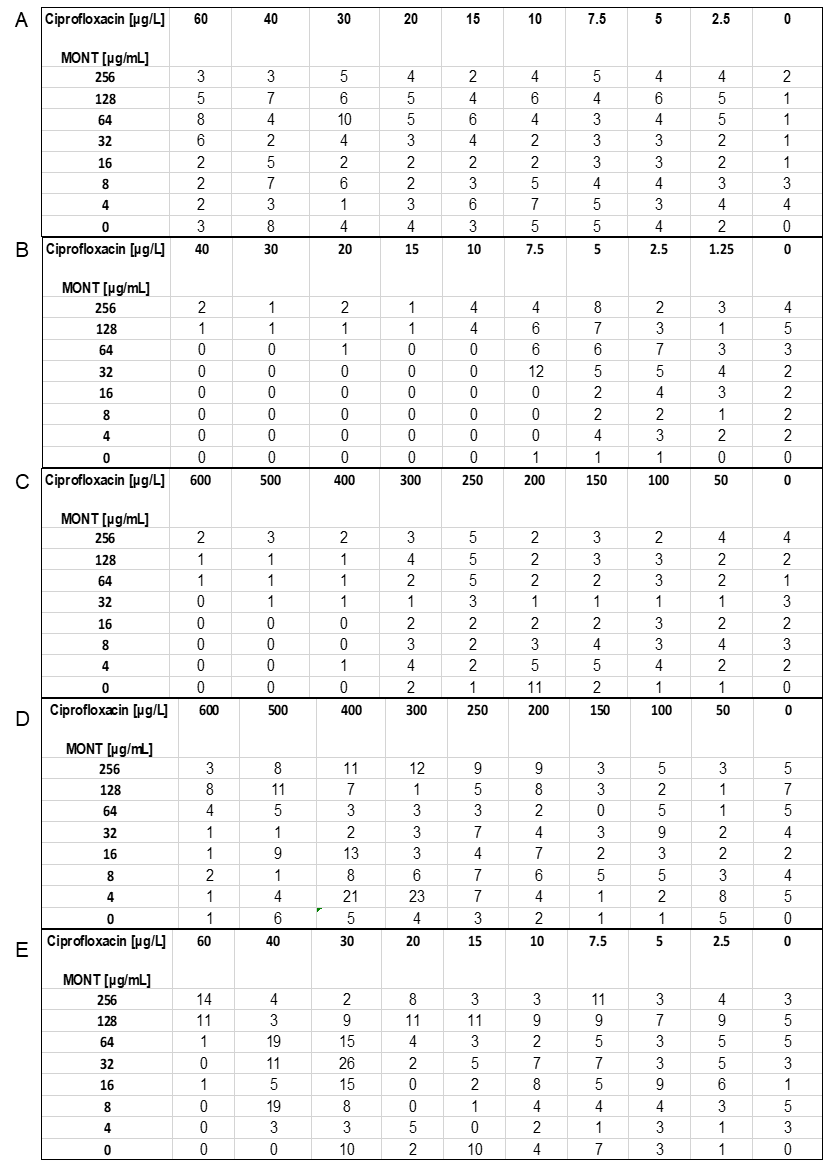

Figure S1: Standard deviations of relative growth [%] shown in Figure 1 of the paper. A. baylyi BD413 (A), E. coli ATCC25922 (B), S. aureus ATCC29213 (C), E. faecalis ATCC29212 (D) and B. subtilis W23 (E).

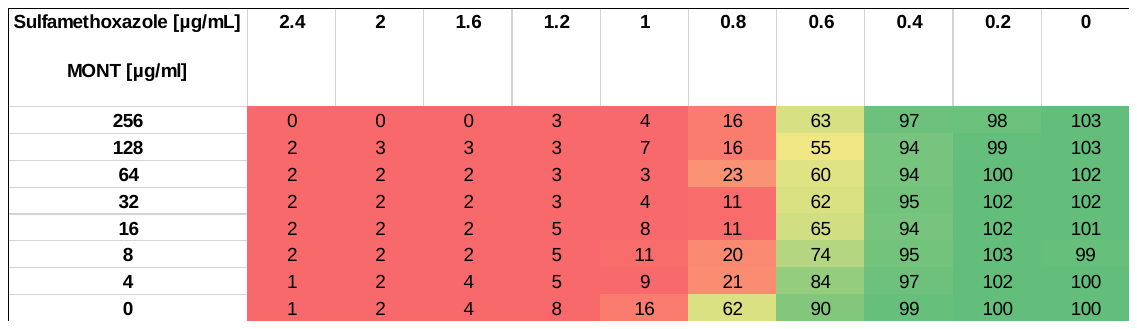

Figure S2: Relative growth [%] of A. baylyi BD413 with different concentrations of SMX and MONT in a 96 well plate. The optical density after 20 h of incubation in each well was compared to growth without antibiotic and colloid and expressed as percentage.

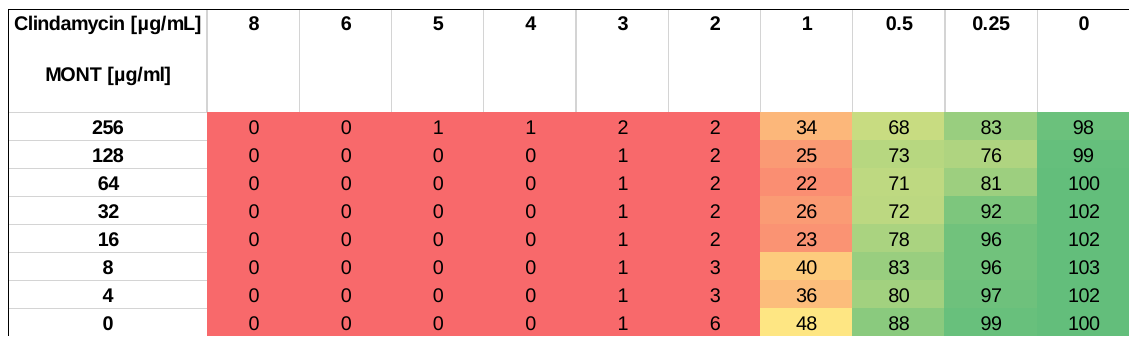

Figure S3: Relative growth [%] of A. baylyi BD413 with different concentrations of CLI and MONT in a 96 well plate. The optical density after 20 h of incubation in each well was compared to growth without antibiotic and colloid and expressed as percentage.

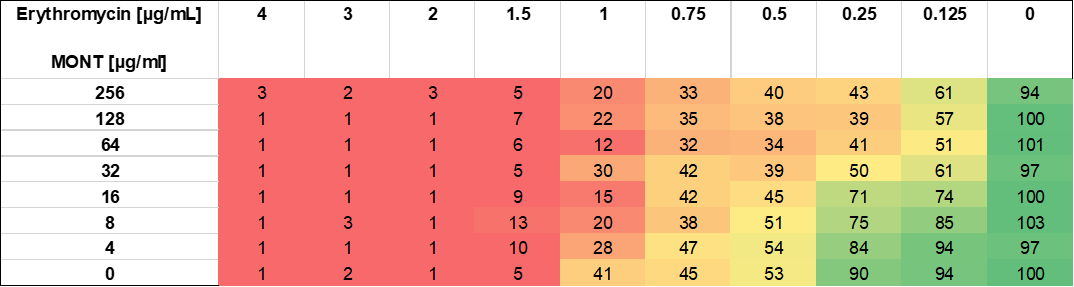

Figure S4: Relative growth [%] of A. baylyi BD413 with different concentrations of ERY and MONT in a 96 well plate. The optical density after 20 h of incubation in each well was compared to growth without antibiotic and colloid and expressed as percentage.

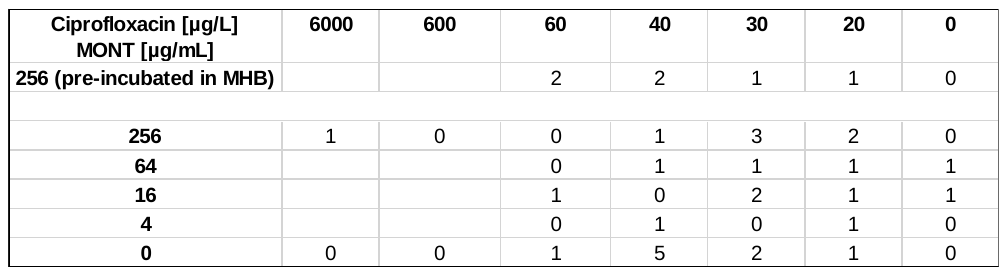

Figure S5: Standard deviations of relative growth [%] shown in Figure 2 of the paper.

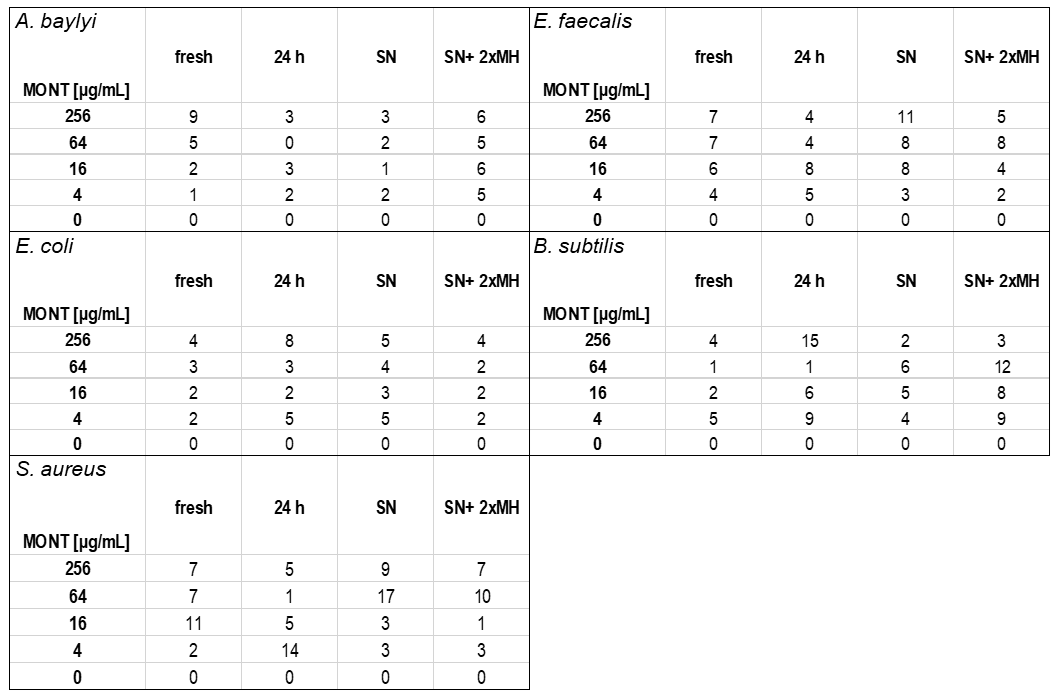

Figure S6: Standard deviations of relative growth [%] shown in Figure 3 of the paper. A. baylyi BD413, E. coli ATCC25922, S. aureus ATCC29213, E. faecalis ATCC29212 and B. subtilis W23.
